# Limited variation in microbial communities across populations of *Macrosteles* leafhoppers (Hemiptera: Cicadellidae)

**DOI:** 10.1101/2024.01.28.577611

**Authors:** Sandra Åhlén Mulio, Agnieszka Zwolińska, Tomasz Klejdysz, Monika Prus-Frankowska, Anna Michalik, Michał Kolasa, Piotr Łukasik

## Abstract

Microbial symbionts play important roles in insect biology, but their diversity, distribution, and dynamics over time across host populations are poorly understood. We surveyed the spatio-temporal distribution of bacterial symbionts in the broadly distributed and economically significant leafhopper genus *Macrosteles*, with emphasis on *Macrosteles laevis*, using host and symbiont marker gene amplicon sequencing. The cytochrome oxidase I (COI) gene data revealed no strong genetic differentiation across *M. laevis* populations, significant levels of heteroplasmy, and multiple cases of parasitoid infections. 16S rRNA data confirmed the universal presence of the ancient nutritional endosymbionts *Sulcia* and *Nasuia* and a high prevalence of *Arsenophonus*. Interestingly, in contrast to most previously surveyed species, in *M. laevis* we found only occasional cases of infection with facultative endosymbionts and other bacteria. There was no significant variation in symbiont prevalence across populations, or among sampling years for the same population. Facultative endosymbionts including *Rickettsia*, *Wolbachia*, *Cardinium*, and *Lariskella*, were more common in other Macrosteles species. Combined, our data demonstrate that not all species show clear spatial and temporal variation in genetic structure and microbial prevalence. However, simultaneous characterization of host and symbiont marker gene amplicons in large insect collections can help understand the dynamics of host-microbe interactions.

## Introduction

Microorganisms are diverse and abundant across all kinds of habitats on Earth, but the microbiota of eukaryotic organisms have attracted particular attention due to their ecological, evolutionary, but also biomedical and agricultural significance (McFall-Ngai *et al*., 2013). These host-microbe associations are diverse taxonomically and functionally, especially in insects: ranging from mutualism, through commensalism, to pathogenicity, they have, in many cases, dramatically influenced the biology and evolution of hosts (Douglas, 2015). In insects, three main symbiont categories are typically distinguished. First, obligate nutritional endosymbionts are strictly heritable, endocellular microbes critical to the survival, function, and reproduction of the insect host as they provide essential amino acids and vitamins lacking in their diet, filling the gaps of the nutritional needs of their host (Baumann, 2005; Moran *et al*., 2008; Sudakaran *et al*., 2017). Due to their indispensable roles and strict maternal transmission, nutritional endosymbionts are expected to occur in all individuals of a species (Baumann, 2005; Moran *et al*., 2008). The second functional category, facultative endosymbionts, are more variable in their distribution across species, populations, and host tissues, as well as in their transmission mechanisms. Although not critical for survival, they may provide important benefits that are often related to defense, or else, manipulate host reproduction (Sudakaran *et al*., 2017). Facultative symbionts are typically transmitted maternally (vertically), but are capable of occasional transmission across host lines or species (horizontally). Thirdly, diverse microorganisms frequently colonize the host gut or body surfaces. Some of these microbes, inducing a wide range of effects, have co-diversified with hosts for a long time, but others form less stable associations (Sudakaran *et al*., 2017). Finally, pathogens - whether attacking a host species or vectored by it - form an important fourth category of microbes that may not conform to the definition of symbiosis as a long-term interaction among organisms but can play important roles in host biology.

The insects in the Hemipteran suborder Auchenorrhyncha are among the best-known symbiotic clades. Feeding on nutrient-imbalanced plant sap, they require nutritional supplementation from symbiotic microbes (Baumann, 2005; Bennett and Moran, 2013). As such, around 300 MYA, the ancestor of Auchenorrhyncha developed an association with a nutritional symbiont, a Bacteroidetes bacterium that we now know as *Candidatus* Sulcia muelleri (hereafter *Sulcia*), which has co-diversified with hosts since then (Moran *et al*., 2005). Additional obligate symbionts, likely acquired independently in different Auchenorrhyncha superfamilies (Deng *et al*., 2023), complemented *Sulcia* in nutrient provisioning, co-diversifying with hosts. When the family Cicadellidae (leafhoppers) separated from treehoppers about 180 MYA, they were accompanied by the obligate endosymbiont known as *Candidatus* Nasuia (hereafter *Nasuia*) (Johnson *et al*., 2018). In the superfamily Fulgoroidea, this second obligate symbiont is *Vidania*, in Cercopoidea *Zinderia* (Bennett and Moran, 2013) and in cicadas, *Hodgkinia* (McCutcheon *et al*., 2009).

While these ancient symbiotic relationships can be stable, symbiont replacement and complementation by other symbionts that provide nutrients have occurred several times throughout evolutionary history in most Auchenorrhyncha groups. For example, in many lineages of cicadas, *Hodgkinia* has been replaced by *Ophiocordyceps* fungi (Matsuura *et al*., 2018). In only rare cases, hosts seem to have lost their symbionts entirely, such as in the largely mesophyll-feeding leafhopper subfamily Typhlocybinae (Moran *et al*., 2005). At the same time, infections with facultative symbionts have often been reported, such as *Arsenophonus* (Michalik *et al*., 2023) where the function is generally unknown despite instances of a nutritional function (Nováková *et al*., 2009), or *Wolbachia* (Qu *et al*., 2013) which has been estimated to occur in about 50% of all insect species (Kaur *et al*., 2021). *Rickettsia* is another widespread symbiont in many insect groups (Pilgrim *et al*., 2021). Despite their common existence, the functions of facultative symbionts are rarely well understood. These secondary symbionts may manipulate reproduction or offer protection against biotic and abiotic challenges, including pathogens, parasites, toxic chemicals, or heat (Oliver *et al*., 2010), Indeed, in species such as the pea aphid, extensive fluctuation in the prevalence of facultative endosymbiont species across populations and seasons was observed, driven to at least some extent by their defensive effects (Smith *et al*., 2015, 2021). In Auchenorrhyncha, little is known about the composition and function of gut microbiota (Michalik *et al*., 2023), despite their importance for many other insects (Engel and Moran, 2013).

Due to their mode of feeding, Auchenorrhyncha may cause feeding damage, but more importantly, vector plant pathogens, such as the agriculturally important genus *Candidatus* Phytoplasma (Mollicutes) (hereafter *Phytoplasma*). These phloem-limited plant parasitic bacteria can cause extensive damage to more than 700 plant species, including maize (Ramos *et al*., 2020), carrot (Clements *et al*., 2021), grapevine (Moussa *et al*., 2023) and many other important crops. Therefore, managing these vector species is of great economic importance even though estimates of economic loss are lacking. More than 75% of the known *Phytoplasma* vectors belong to the leafhopper subfamily Deltocephalinae (Weintraub *et al*., 2019). This group includes the leafhopper genus *Macrosteles*, a widespread and morphologically elusive genus with around 90 species described to date, several of which are known to vector phytoplasmas (Frost *et al*., 2011; Weintraub *et al*., 2019).

The association of *Macrosteles* spp. with ancient nutritional endosymbionts *Sulcia* and *Nasuia* have been comprehensively characterised, with *M. quadrilineatus* as one of the model systems for the study of the mechanisms of these interactions (Bennett *et al*., 2016; Mao and Bennett, 2020). Symbiotic associations were also surveyed in *M. striifrons* and *M. sexnotatus* from Japan where *Sulcia* and *Nasuia* were reported alongside facultative symbionts *Rickettsia* and *Wolbachia*, and putative gut bacteria *Burkholderia*, *Pantoea*, and others (Ishii *et al*., 2013). Further, *M. laevis*, characterised using microscopy and sequencing-based techniques, in addition to *Sulcia* and *Nasuia* was shown to harbour the symbiont *Arsenophonus* in a particularly unusual location: within the cytoplasm of *Sulcia* cells (Kobiałka *et al*., 2016). But while we know the identity of some of *Macrosteles* symbionts, little is known about the diversity of their microbiota and its spatio-temporal variation across species and populations.

Given the lack of such research combined with the economic incentive of surveying a vector species, the goal of this study was to comprehensively survey and characterise spatio-temporal patterns of microbial diversity and distribution in Polish and European populations of *M. laevis* and a few other species in the genus *Macrosteles*. We aimed, firstly, to test the utility of COI amplicon sequencing and barcoding (short standardised DNA sequences used to identify species (Chac and Thinh, 2023)) as a means of delimiting host species, verifying their identity, and detecting parasites or parasitoids, as potential explanatory variables of microbiome composition. Secondly, we wanted to explore the genetic diversity among the host species and populations, asking whether the intraspecific genetic structure could explain the symbiont diversity patterns. Thirdly, we aimed to characterise the diversity and distribution of *Macrosteles* symbionts and the variation in *M. laevis* microbiota across populations and over time. Fourthly, we wanted to investigate potential patterns of *Phytoplasma* infection and its correlation with the presence of other microbes. We addressed these goals by sequencing insect and bacterial marker gene amplicons for the collection of over 300 specimens from different populations.

## Materials and methods

### Specimens acquisition and identification

The majority of specimens were collected using sweep netting in Poland between 2015-2020, and additional samples were collected or obtained from the UK, USA and Sweden (Supplementary Tables S1-S2). Specimens, identified to species level based on morphological features, were stored in 95% ethanol in a freezer until processing. Species identifications included *M. laevis*, *M. sexnotatus*, *M. cristatus*, *M. maculosus*, *M. viridigreseus* and *M. quadrilineatus*, but *M. laevis* was by far the most dominant. In total, 371 specimens were processed, and for 307 we obtained reliable data and included them in this study (as explained below). A substantial portion of the samples included 157 individuals of *M. laevis*, collected from two distinct Polish agroecosystems: one in Szczecinek and the other in Sośnicowice, spanning the years 2015 to 2018, with 40 insects collected each year. These specimens were obtained from plots adjacent to oilseed rape fields, which primarily comprised wild plants, predominantly grasses.

### DNA preparation and extraction

The DNA from the collection that included 157 samples collected in Szczecinek and Sośnicowice before 2020 was extracted using the NucleoSpin Tissue kit (Mecherey-Nagel). Whole specimens were placed in tubes filled with a lysis buffer provided by the manufacturer and homogenised using a mini pestle. For further steps, the manufacturer protocol was followed. For amplicon sequencing, we selected samples that tested positive and negative for *Phytoplasma* infections in the earlier nested-PCR screen, which followed the standard method for phytoplasma detection as previously described (Zwolińska and Borodynko□Filas, 2021). Our goal was to include 20 samples from each of the two categories — positive and negative — for each site-date combination, although there were instances where lower numbers were available.

21 samples from Poland, Kampus UJ, were processed using three different extraction approaches: DNeasy PowerSoil (Qiagen) and DNeasy Blood and Tissue kit (Qiagen), for which the manufacturer protocols were followed, and a custom protocol, described below. For each of the three kits, seven specimens were processed, four as whole specimens and with only the abdomens used for the remaining three. As we found no consistent differences among the protocols, the remaining 193 samples were processed using whole specimens and our custom protocol.

Specimens were each placed in 2 ml screw-cap tubes together with a lysis buffer (GitHub repository: https://github.com/SymSandra/Macrosteles-project/), proteinase K and ceramic beads. Samples were then homogenised in the homogenizer for 30 s at 5 m/s two times, and incubated in a thermal block at 56 degrees Celsius for 2 hours. 20% of the homogenate for each sample was used for DNA purification using SPRI magnetic beads (Sera-mag, SpeedBeads magnetic carboxylate modified particles, Cytiva). Remaining product was stored in a freezer.

To track possible contamination in samples, negative controls were used in the form of water for both extraction and subsequent steps of PCR and indexing, amounting to 24 control samples in total for all 371 samples processed.

### Amplicon library preparation and sequencing

During the first step, we targeted two regions - insect COI, using primers BF3 and BR2 (Elbrecht and Leese, 2017) and hypervariable regions V1-V2 and V4 of bacterial 16S rRNA gene, using primers 27F and 338R (Walker *et al*., 2015) and 515F and 806R (Apprill *et al*., 2015; Parada *et al*., 2016), respectively. These primer pairs, with partial Illumina adapters at the 5′ end, were combined in a single multiplex reaction with Qiagen Multiplex DNA polymerase. Then, after purification using SPRI magnetic beads, PCR products were used as templates for the second, indexing PCR, with a different set of uniquely labelled oligonucleotides used as primers for each sample. Subsequently, PCR products were verified and roughly quantified using a 2.5% agarose gel stained with Midori Green. Libraries were then pooled based on gel band intensity, cleaned with Promega magnetic beads, and sequenced in a partial Illumina MiSeq V3 lane (2 × 300 bp reads) at the Institute of Environmental Sciences of Jagiellonian University. All protocols are provided and described in the following GitHub repository: https://github.com/SymSandra/Macrosteles-project/.

### Amplicon data analysis

Amplicon libraries were processed following a pipeline detailed at a dedicated GitHub page (https://github.com/SymSandra/Macrosteles-project/), closely resembling that described in recent publications (Kolasa *et al*., 2023; Valdivia *et al*., 2023). Briefly, reads were divided into bins corresponding to the targeted marker regions based on primer sequences. Then, each bin was analysed separately using custom protocols combining vsearch, usearch, and custom steps. Forward and reverse reads were assembled into contigs and quality-filtered. Contigs were dereplicated, and representative sequences were chosen, with singleton sequences removed from further analysis. Samples were denoised with usearch and aligned against a customized reference database based on MIDORI (Leray *et al*., 2018) for COI and SILVA 138 (Quast *et al*., 2012) for 16S rRNA. For analysis of V1-V2 and V4 16S rRNA gene sequences, we used UCHIME (Edgar *et al*., 2011) to screen and remove chimeric sequences prior to taxonomy assignment. Finally, the sequences were clustered at 97% identity level, and OTU (Operational Taxonomic Unit) and zOTU (zero-radius OTU) tables were produced. For COI data, we chose the most abundant sequence as the barcode for each sample.

Finally, we used a decontamination procedure for 16S rRNA samples, using as references multiple negative control samples for different steps of library preparation (DNA extraction, first PCR, and a second PCR with indexing). The custom Python script recognizes and filters out OTUs assigned based on the comparison of the relative abundance of genotypes among negative controls and samples, as well as removing OTUs assigned as mitochondria, chloroplasts, Eukaryotes, or Archaea (for more details, see the GitHub repository). Resulting data tables were manually filtered so that only the zOTUs represented by at least 100 reads and representing at least 5% in at least one sample, were specifically considered in distribution analyses, with the remainder classified as “Others”. Samples with less than 100 COI reads representing the barcode (that is, the dominant *Macrosteles* genotype), or with less than 1000 16S-V4 reads, were excluded from all comparisons.

### Interpretation and visualisation

A representative sequence for each COI zOTU was used to create a phylogenetic tree in MEGAX (Kumar *et al*., 2018; Stecher *et al*., 2020), using the Maximum Likelihood approach with 1000 bootstraps. Heatmap for 16S rRNA representative sequences and boxplots were created using Rstudio, version 1.2.5033 and packages: usethis (Wickham *et al*., 2022) devtools (Wickham *et al*., 2022), pheatmap (Kolde, 2019), RColorBrewer (Neuwirth, 2022) and gridBase (Murrell, 2014). The map and figures were adjusted using Inkscape, version 1.1.2. Barcharts and Chi-square tests were made with Excel, version 16.59. Host population circles were created using Processing software, version 3.5.4.

## Results

### Amplicon sequencing data overview

Out of 371 insects initially processed for this study, 65 were excluded because of an insufficient number of reads for either the COI region (≧100), preventing molecular identification, or for the V4 region of the 16S rRNA (≧1000). Most of the discarded specimens represented the Swedish collection, stored under suboptimal conditions which led to greater DNA degradation.

Consequently, 306 libraries were used for core analyses combining COI and V4 data.

The amount of data for the V1-V2 region of the 16S rRNA gene was generally less than for V4, and 77 additional libraries were further excluded from the comparison among these two gene regions. V1-V2 data were not used for other comparisons.

### COI-based host species delimitation and genotype diversity

Across 306 samples, we identified 249 COI genotypes (zOTUs), clustering into 43 OTUs. 44 of these genotypes, representing a large majority of reads (90.8%) were clustered into 7 OTUs which were taxonomically annotated as one of seven *Mactosteles* spp. and considered as representing mitochondrial COI sequences, or barcodes (Supplementary Table S3-4). We identified another group of OTUs also classified as *Macrosteles*, but always accompanying the “barcode” OTUs and not exceeding 10% of the total number of reads in the library, but generally closer to 1%. We assume that these secondary OTUs represent nuclear pseudogenes (numts), or other biologically inactive sequences. However, a curious case is OTU10, comprising between 0 and 6% reads in all *M. laevis* individuals except for one, SWEAM3, where it makes up 75%, dominating over the “barcode” OTU1. We are not sure whether this represents a true biological phenomenon, or a laboratory artifact, for example, related to the patterns of DNA degradation.

For all but 10 specimens, COI sequence-based taxonomic annotations matched morphology-based IDs, but in the remaining cases, the barcode sequences matched different *Macrosteles* species than expected. Specifically, five out of six specimens previously identified as *M. sexnotatus* and one specimen of *M. cristatus* had barcodes matching *M. laevis*. One specimen identified as *M. laevis* belonged to the species *M. cristatus*. One specimen of *M. cristatus* matched the barcode of *M. viridigreseus.* Lastly, two specimens morphologically identified as *M. laevis* had the closest match to *M. frontalis* after a Blast search. Given the known challenges with morphology-based identifications of this elusive genus, the results of barcoding were used for further analysis.

The analysis of intra-OTU COI genotype diversity provided interesting insights into *Macrosteles* population structure. In *M. laevis*, one genotype - a consensus sequence for all genotypes in the OTU - dominated the profiles of most specimens from all locations. However, in 43 out of 270 (15.9%) *M. laevis* individuals from across populations and sampling years, we identified other genotypes that contributed >5% of the total OTU, and usually co-ocurred with the consensus one. To identify sequence variants that are biologically realistic, rather than representing sequencing artifacts, for the characterization of genotype-level patterns we only considered genotypes represented by ≧100 reads and with a relative abundance threshold of ≧5% of at least one library. For the most abundant species, *M. laevis*, where the barcode was represented by OTU1, 27 out of 60 zOTUs in that OTU exceeded this relative abundance threshold (Fig. 1). Almost all of these genotypes differed at only a single nucleotide position from the dominant one, but their distribution showed no consistent geographic or temporal patterns (Figure 1). We interpret these patterns as heteroplasmy - the presence of more than one mitochondrial genotype within an individual (Avise, 2000), likely resulting from recent point mutations that have spread but not yet reached fixation within individuals or populations. Likewise, we found more than one COI genotype in *M. viridigreseus*, with three equally abundant genotypes in the British population. The single Polish specimen of *M. viridigreseus* had a distinct genotype to those occurring in the British population. In *M. quadrilineatus*, two genotypes were found, with one genotype clearly dominant across the samples but with two specimens displaying a second genotype.

**Figure 1.**
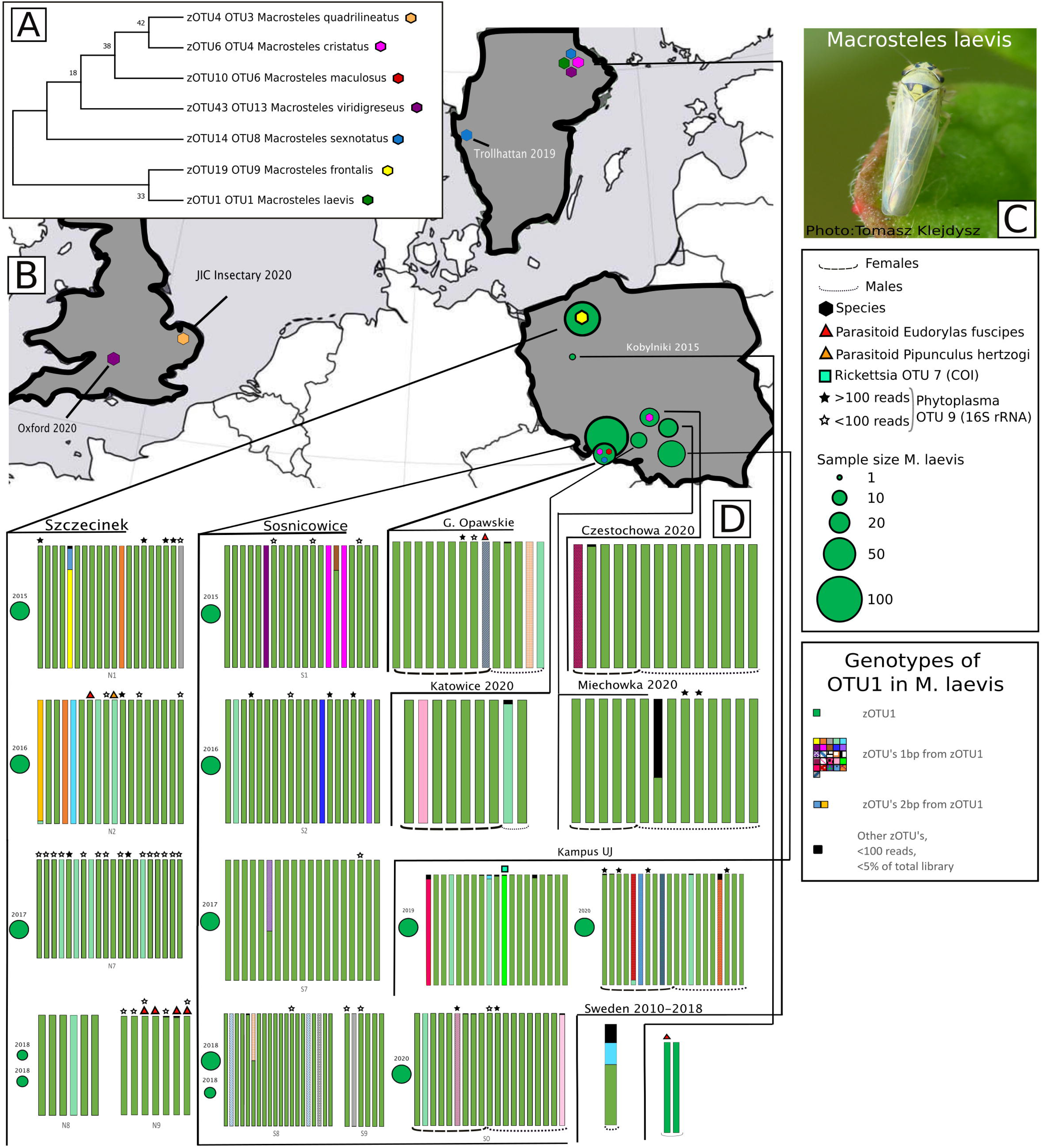
**A**, Maximum likelihood tree for the sampled Macrosteles species based on the dominant zOTU sequence from each respective OTU. Support values are based on 1000 bootstraps. **B**, Sample locations for all specimens. Per-site sampling effort shown for *Macrosteles laevis*, with green circle size representing the number of specimens collected across all seasons. Hexagon colours correspond to species as in panel A. **C**, Photo of *M. laevis* by Tomasz Klejdysz. **D**, COI genotype diversity of *M. laevis* (OTU1) as relative abundance of zOTUs in percentage based on the cutoffs of >100 reads and making up at least 5% of the library. Each bar corresponds to one specimen. Size dependent circles correspond to sampling effort across different years for sites with multiple sampling seasons.

### COI amplicon data inform about parasitoid, *Wolbachia*, and *Rickettsia* infections

Interestingly, a significant share of reads in our COI dataset did not match leafhoppers, instead providing insights into their interactions with other organisms. Two OTUs matched NCBI database records of *Eudorylas fuscipes* and *Pipunculus omissinervis* (both Diptera: Pipunculidae) with 100% and 98.8% identity, respectively. The first (OTU2, *E. fuscipes*) was represented by four genotypes distributed across 11 individuals, including 2 where it was represented by >100 reads. The second (OTU16, *P. omissinervis*) occurred in a single specimen only. Species of the family Pipunculidae are known parasitoids of Auchenorrhyncha and *E. fuscipes* is known to parasitize *M. laevis* (Bańkowska, 1989). Both parasitoids occurred only in Polish specimens of *M. laevis* across several years.

Another set of COI OTUs matched *Rickettsiales* bacteria *Wolbachia* and *Rickettsia*, known as facultative endosymbionts in a wide variety of insects. *Wolbachia* OTU5 was present in 2 specimens of *M. viridigreseus*, one specimen of *M. laevis*, and 1 specimen of *M. maculosus*, and OTU22 - in two specimens of *M. laevis. Rickettsia* was represented by two COI OTUs, each in a distinct *M. laevis* individual.

### Microbiota diversity comparison based on two 16S rRNA regions

Across 229 libraries with at least 1000 reads for both V4 and V1-V2 regions of 16S rRNA gene (Tables S5-8), we compared the presence of more abundant clades, focusing on OTUs that exceeded 1% relative abundance in at least one library. For the V4 region, we identified 21 such OTUs, comprising 99.88% of reads in a library on average. For the V1-V2 region, we identified 29 such OTUs, comprising 99.83% of reads in a library on average (Supplementary Table S9).

Both datasets were dominated by the OTUs of heritable endosymbionts. *Sulcia* OTU and *Arsenophonus* OTUs were abundant in both datasets, although in the latter case, differential clustering of genotypes into OTUs was particularly evident (Figure 2). On the other hand, *Nasuia* was well-represented in only the V4 dataset - identified in every library and comprising 30.92% of reads on average, whereas in the V1-V2 dataset, it was only found in 51 libraries, at low abundance. Facultative endosymbionts *Cardinium*, *Rickettsia*, *Wolbachia*, and unclassified Gammaproteobacterium were better represented in the V1-V2 dataset than in the V4 dataset, as was *Phytoplasma* (Figure 2). However, the six-fold greater V4 sequencing depth (on average) may explain why *Phytoplasma* was detected in fewer V1-V2 than V4 libraries (24 vs 48).

**Figure 2.**
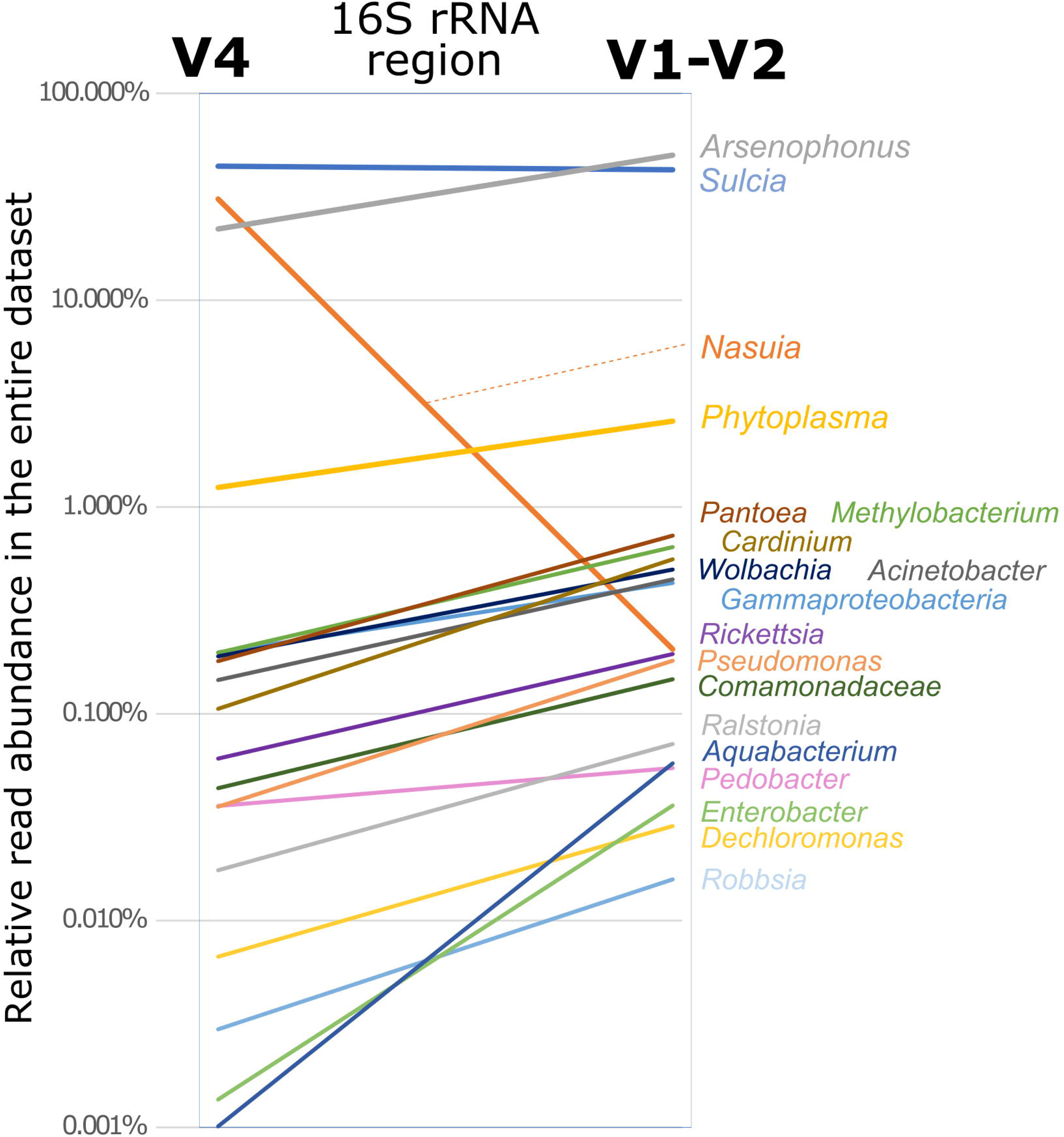
The comparison of averaged relative abundances of dominant bacterial genera among amplicon datasets for 16S rRNA V4 and V1-V2 regions for the same 229 *Macrosteles* samples.

The lack of overlap among the 16S rRNA regions has complicated direct OTU-to-OTU comparisons among less abundant bacteria, and we based further comparisons primarily at genus-level classification (Table S9), in some case also using full-length 16S rRNA sequences from Genbank to establish equivalence. We concluded that all genera represented by OTUs that exceeded 1% in at least one V4 library were represented among the more abundant OTUs in the V1-V2 dataset. Likewise, the bacterial genera that exceeded 1% relative abundance in at least one V1-V2 library had equivalents among in the V4 dataset.

We conclude that with the exception of *Nasuia*, the V1-V2 and V4 regions of 16S rRNA gene identify overlapping bacterial clades and reconstruct similar distributions. Many less abundant genera have higher representation in the V1-V2 dataset, which could have facilitated their detection in the case of even sequencing coverage. However, because of the much lower overall read numbers for the V1-V2 region, we have decided to rely on V4 region data only for the subsequent analyses.

### The endosymbiont diversity across *Macrosteles* species

The microbial communities of the surveyed *Macrosteles* leafhoppers were dominated by the previously reported heritable endosymbionts (Figure 3). All specimens hosted the obligate nutritional symbionts *Sulcia* and *Nasuia*. These two bacteria dominated the microbiota in the vast majority of specimens, with *Sulcia* making up 45.3% of the microbial community on average (range 4.12-84%), and *Nasuia* 26.6% on average (range 1.95-80%). All characterised *Macrosteles* leafhoppers, regardless of the species, hosted the same *Sulcia* genotype. *Nasuia* was represented by ten genotypes grouped into two OTUs, with each species hosting a distinct genotype, or a set of genotypes. Some *M. laevis specimens* harbored a distinct genotype from the dominant one for the species, with a single (zOTU038) or two-nucleotide difference (zOTU033 and zOTU041).

**Figure 3.**
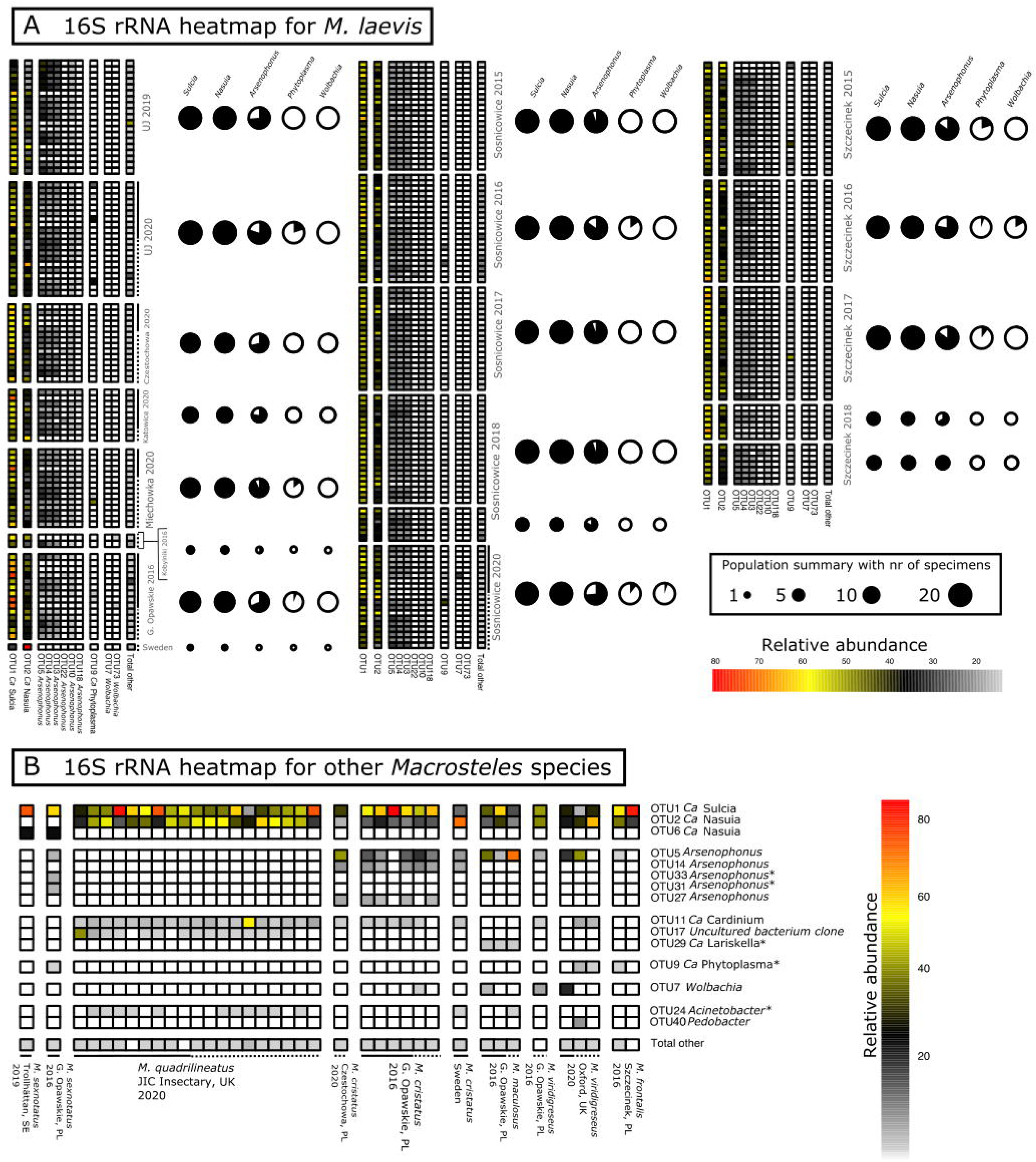
**A**, Relative abundance of the dominant bacterial 97% OTUs across Polish populations of *M. laevis*, based on amplicon data for the V4 region of 16S rRNA Rows correspond to specimens, and columns - to bacterialOTUs. Column “Total other” contains the sum of all other OTU’s present in each specimen. White squares equals 0% relative abundance. For all sampled populations, circle size represents the sampling depth for the population, and the black area of the circle - the proportions of individuals infected with the particular symbiont. **B**, Symbiont relative abundance across populations of other *Macrosteles* species. Each row corresponds to one bacterial 16S rRNA OTU and each column corresponds to one specimen. Row “Total other” contains the sum of all other OTU’s present in each specimen. OTUs marked with an asterisk signifies OTU’s that did not pass the cutoff criteria but were selected to highlight distribution. Full line corresponds to females and dotted line to males.

We also identified OTUs matching known facultative endosymbionts of insects in the majority of studied individuals. Most notably, OTUs identified as *Arsenophonus,* a Gammaproteobacterium previously shown to inhabit the cells of *Sulcia* in one studied population of *M. laevis* (Kobiałka *et al*., 2016) occurred in 222 of 270 (82.2%) *M. laevis* specimens, with relative abundance within libraries ranging from 0.004%-68.8%. In *M. laevis*, we detected four OTUs classified to the genus *Arsenophonus* that were consistently present in infected specimens. Because of the consistent abundance of these OTUs relative to each other and the high degree of sequence divergence among rRNA operons within the genomes of previously characterised strains (Nováková *et al*., 2016), we concluded that the different OTUs present in single specimens almost certainly represent a single microbial strain. The *Arsenophonus* prevalence within populations ranged from X to Y%, with no consistent differences across sites or sampling dates. The same pattern of several co-occurring OTUs was present in *M. cristatus* and *M. sexnotatus*. However, In other *Macrosteles* species, *Arsenophonus* was generally represented by different OTU combinations and made up less than 10% on average.

Facultative endosymbiont *Wolbachia*, represented by two OTUs, was relatively rare and found only in a small proportion of specimens. The OTU007 of *Wolbachia* was present in two specimens of *M. viridigreseus,* one specimen of *M. laevis,* one specimen of *M. maculosus* and one specimen of *M. cristatus,* from the Polish population. Notably, in four out of these five specimens, Wolbachia was also detected in COI amplicon data. *Wolbachia* OTU073 was present in nine *M. laevis* specimens, and its relative abundance did not exceed 0.77%. Notably, five of these specimens were among the twelve that contained parasitoid *Eudorylas fuscipes* COI reads. It seems plausible that the symbiont actually infected the parasitoid rather than the leafhopper, but the low relative amounts of parasitoid tissue hindered their reliable co-detection.

In the V4 dataset, we also found *Trichorickettia* in two *M. laevis* specimens, each from a different Polish population. Further, most individuals of *M. quadrilineatus* were infected by a bacterial OTU17, likely a Gammaproteobacterium, but without close similarity to any named bacteria in the NCBI database.

We also discovered two putative facultative endosymbionts not previously reported in *Macrosteles* leafhoppers. Firstly, we found *Cardinium* in all *M. quadrilineatus* and in seven out of eight *M. cristatus* from Poland and one from Sweden. It was also found in two of the British *M. viridigreseus* specimens. The one specimen of *M. viridigreseus* missing the symbiont *Cardinium* was infected by *Wolbachia*. Finally, we identified an Alphaproteobacterium classified as *Lariskella* in three specimens of *M. maculosus* with a relative abundance ranging from 0.9-1.17%. Combined, the seven symbionts listed above made up 97.2% of reads in a library on average, and together with *Phytoplasma* described in the next section, 98.2%

The other microbes were less common and abundant in the dataset. They included *Enterobacterales* genus *Pantoea, Rhizobiales genus Methylobacterium, Pseudomonadales* genera *Acinetobacter* and *Pseudomonas*, *Burkholderiales* genera *Massilia*, *Acidovorax*, *Ralstonia*, *Janthinobacterium*, *Dechloromonas*, and *Robbsia*, *Sphingobacterales* genus *Pedobacter*, and others. Of these, *Pantoea* OTU019 was relatively most widespread and abundant, infecting 25.8% of all specimens, including 27% of specimens in *M. laevis* and 10.5% in *M. quadrilineatus,* 37.5% in *M. cristatus* and 33.3% in *M. maculosus*. It made up an average of 0.2% of the microbial community (max. 14.05%). Other OTUs were less prevalent and abundant, and none of them showed consistent associations or distribution patterns across species and populations that we would expect in cases of a specific and biologically significant association (Supplementary Table S5-6).

### Surveying *Phytoplasma* prevalence across *Macrosteles* individuals

The plant-pathogenic bacterium *Phytoplasma* was detected in 57 individuals, but was generally found at relatively low abundance: in one individual, it was 44.8% and in a few others ∼10%, but typically, it was well below 1% (Figure 4a, Supplementary table S5-6). This large variation among individual insects in *Phytoplasma* relative abundance may explain the limited overlap between its detection based on amplicons as opposed to diagnostic nested PCR (Zwolińska and Borodynko□Filas, 2021). When looking at the 156 *M. laevis* individuals from Szczecinek and Sośnicowice (Poland) that were pre-selected for microbiome characterization based on the results of *Phytoplasma* screen, we found a discrepancy between *Phytoplasma* detection based on nested PCR and amplicon data (Figure 4A). Out of 59 specimens selected as *Phytoplasma*-positive based on nested PCR, in only 24, we found 16S rRNA reads representing this bacterium. At the same time, out of 97 specimens selected as *Phytoplasma*-negative based on nested PCR, we found 16S rRNA reads matching this microbe in 18 samples. While *Phytoplasma* was more likely to be detected in amplicon data for nested PCR-positive specimens, 16S rRNA amplicon sequencing on its own appears insufficient as a means of detecting *Phytoplasma* presence. Outside of these pre-screened populations, the remaining occurrences of *Phytoplasma* were found in 11 *M. laevis* from other populations, single specimens of *M. frontalis* and *M. sexnotatus*, and two specimens of *M. viridigreseus*.

**Figure 4.**
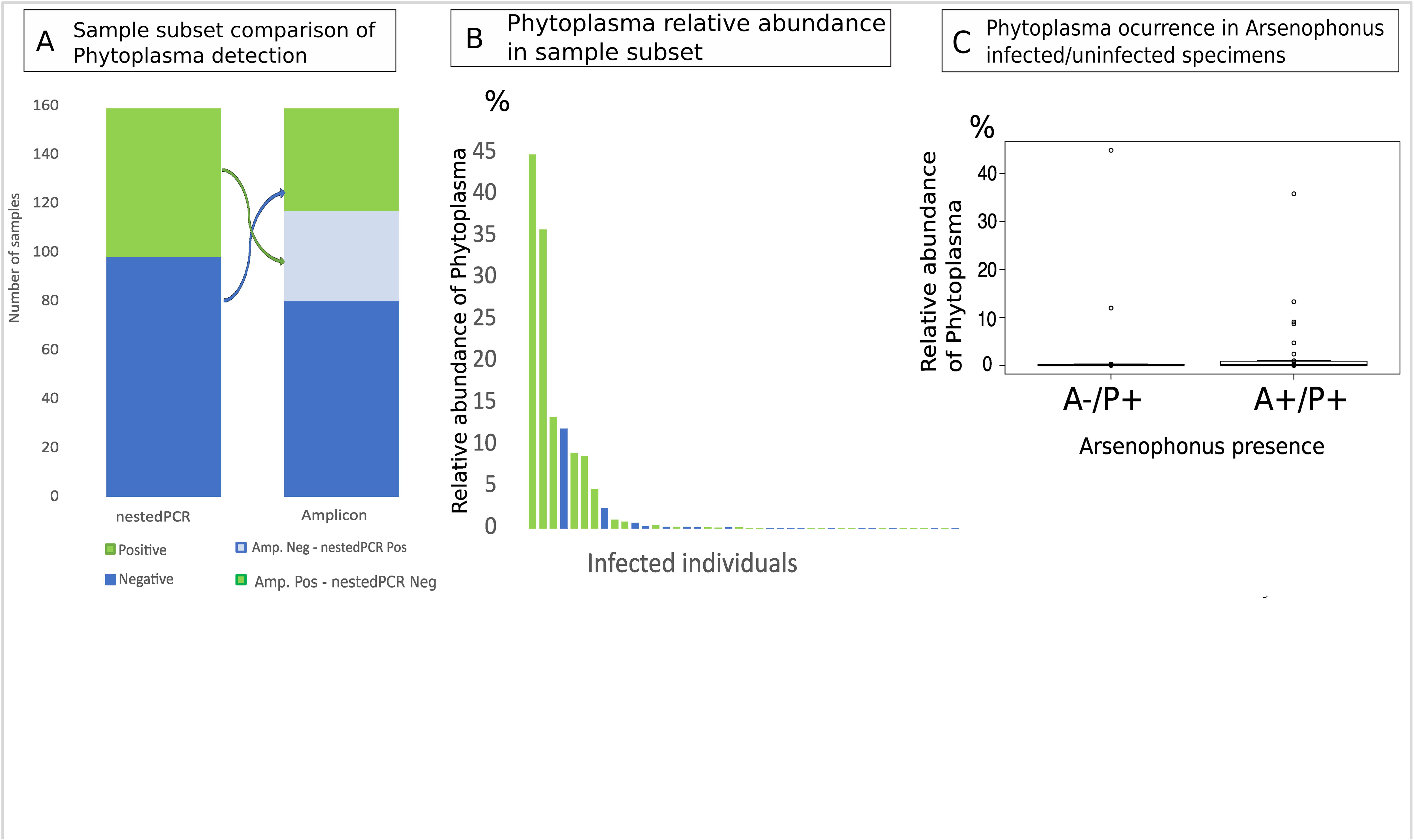
**A,** The comparison in *Phytoplasma* detection efficiency among *M. laevis* individuals from Szczecinek and Sośnicowice that had previously been scored as *Phytoplasma*-positive or negative based on nested PCR. **B,** Relative abundance of *Phytoplasma* in microbial communities of 42 pre-selected infected specimens from Szczecinek and Sośnicowice, based on amplicon sequencing. **C**, Relative abundance of *Phytoplasma* in microbial communities of individuals where *Arsenophonus* was present vs absent

At the genotype-level, *Phytoplasma* was represented by thirteen zOTUs (Supplementary Table S6). The dominant genotype (zOTU14) was found in 35 *M. laevis* specimens from Szczecinek-Sośnicowice collections and was also detected in three additional Polish populations. Two other genotypes, zOTU91 and zOTU100, co-occurred together in all six specimens from four Polish populations of *M. laevis* at a near 50/50 ratio, suggestive of them representing distinct rRNA sequence variants within a single genome.

Further genotypes, zOTU230 and zOTU423, only occurred in specimens from Szczecinek sampled in 2017, and six of seven specimens were co-infected with the dominant genotype (zOTU14). Additionally, zOTU451 only occurred in two specimens from Sośnicowice from 2016. The remaining lower-abundance genotypes occurred in Szczecinek-Sośnicowice collection in a single specimen. Six of the low abundance genotypes occurred in only one specimen. Besides *M. laevis, Phytoplasma* (zOTU14) only occurred in one specimen of *M. sexnotatus*, one specimen of *M. frontalis,* and two specimens of *M. viridigreseus* in which one had an additional genotype (zOTU91). The genotype zOTU137 was only present in two specimens from G. Opawskie.

We also asked whether the *Phytoplasma* abundance/presence correlates with the presence of other microbes in the Szczecinek-Sośnicowice collection. We looked specifically at the correlation with *Arsenophonus* as other microbes were too uncommon. We did not find a correlation between *Arsenophonus* presence or absence and *Phytoplasma* presence or absence across samples of *M. laevis* from Sośnicowice and Szczecinek (Figure 4) (Chi-square test: □2 = 0.53, df = 1, p < 0.46 for all *Phytoplasma* reads and □2 = 0.04, df = 1, p < 0.83).

## Discussion

Our microbiome surveys across seven species of *Macrosteles* leafhoppers revealed the conserved nature of their ancient nutritional symbioses with *Sulcia* and *Nasuia*, and substantial variation among species in the presence of other symbiotic bacteria (Ishii *et al*., 2013). At the same time, the study across 270 individuals of *Macrosteles laevis*, representing several populations and sampling points in Poland and beyond, showed no discernable patterns in the prevalence of other symbionts among or within populations. In both cases, the simultaneous characterization of host and symbiont marker gene data was instrumental for describing and characterising the patterns. Despite the growing popularity of insect COI gene sequencing, or barcoding, as a part of biodiversity surveying efforts (Srivathsan *et al*., 2023), this approach has rarely been incorporated into the study of host-associated microbiota. Here, COI amplicon data provided several pieces of information needed to interpret microbiome data.

First, morphology-based identification of wild-caught insects can be challenging, but the use of COI data can clarify host identity. In our dataset, taxonomically annotated COI data did reveal some misidentifications, supporting the view of the genus *Macrosteles* as an elusive group with a high level of morphological identification difficulty. Most identification is based on males since females often lack distinguishing characteristics (Holzinger *et al*., 2003). However, molecular data proved useful in clearing up taxonomic mistakes and ambiguities.

Second, the composition of the microbial community, and especially heritable symbioses, is known to often correlate with the host phylogeny. COI amplicon data can provide us with at least some insight into intra-species genetic diversity and population history. In a recent broad geographic survey across populations of the spittlebug *Philaenus spumarius*, COI barcodes highlighted the divergence among major clades and revealed smaller-scale genetic variation among populations, which often correlated with symbiont associations (Kolasa *et al*., 2023). In *M. laevis*, no comparable patterns were found. Instead, the identification of the single dominant genotype accompanied by a cloud of genotypes 1-2 bases apart, with no consistent trends or differences among populations, suggested their relatively recent shared ancestry and ongoing diversification. The discovery of substantial levels of heteroplasmy, generally including the dominant COI variant plus one of the variants one base apart, further supports that interpretation. In *M. laevis*, the discovery of that intra-species COI diversity did not help to explain the patterns of microbial associations. However, it would be fascinating to explore its effects in other insect clades where such diversity and heteroplasmy was reported, including neotropical ants (Meza-Lázaro *et al*., 2018), thrips (Frey and Frey, 2004), the honeybee (*Apis cerana*) (Songram *et al*., 2006) and periodical cicadas (Fontaine *et al*., 2007) among others.

Third, COI data for individual insects can provide valuable information about parasitoid or parasite infections. These natural enemies, known to strongly influence insect population dynamics (Abram *et al*., 2019), are likely to alter microbiome profiles - whether by selecting hosts with specific microbiota, shifting the microbial community of the attacked host, or contributing parasites’ microbes to the community profile (Dicke *et al*., 2020). Indeed, in *P. spumarius*, one of *Wolbachia* OTUs only occurred in specimens infected by its specialised parasitoid, suggesting the parasitoid as the source of *Wolbachia* (Kolasa *et al*., 2023). Likewise, known Auchenorrhyncha parasitoids *Pipunculus omissinervis* and *Eudorylas fuscipes* were detected in some *Macrosteles* specimens. Several of those carrying E. fuscipes also contained a *Wolbachia* OTU rare in other samples, suggesting the parasitoid as its source. The imperfect correlation between the reconstruction of parasitoid and symbiont presence may be highlighting the limits of their amplicon-based detection. However, COI data can apparently provide additional information on the presence of alphaproteobacterial symbionts *Wolbachia* and *Rickettsia*, confirming their 16S rRNA-based detection and improving phylogenetic resolution. In our dataset, we found a high correlation between the presence of these microbes in COI and 16S rRNA datasets for the same insects, and we are now working on formally comparing their detection capacity across a diverse range of insects.

The comparison of microbial community reconstructions based on two targeted 16S rRNA regions concluded that the V1-V2 region data was redundant, not contributing additional information relative to the V4 region data while failing to capture the signal of *Nasuia*. Overall, the microbiome data emphasised the stability of ancient nutritional heritable symbioses, previously demonstrated in many clades of Auchenorrhyncha, but not universal across their phylogenetic tree (Bennett and Moran, 2015), and rarely explored as extensively within species. The lack of within-species variation in *Sulcia* and *Nasuia* 16S rRNA sequences further argues for limited divergence among the sampled populations. On the other hand, the slow rate of *Sulcia* genome evolution and limited differences among species within the genus had also been found in other Auchenorrhyncha clades (Campbell *et al*., 2015; Kolasa *et al*., 2023; Michalik *et al*., 2023).

In the genus *Macrosteles*, as in many other hemipteran groups, ancient symbionts are accompanied by accessory symbionts, especially Gammaproteobacteria. In several systems studied to date, they complement ancient symbionts’ biosynthetic pathways and contribute additional nutrients, especially B vitamins (McCutcheon *et al*., 2019). In *M. laevis*, we found *Arsenophonus* across all *M. laevis* populations sampled in Poland, but in neither it has reached fixation, arguing against its essential role despite the very specific and unusual endobacterial localization (Kobiałka *et al*., 2016). *Arsenophonus* was also found in most other *Macrosteles* species except *M. quadrilineatus* - although it has been shown to occur in wild populations of that species too (Bennett and Moran, 2013). We were initially surprised by the apparent diversity of *Arsenophonus*, represented by several OTUs across all sampled *M. laevis* individuals. However, *Arsenophonus* is notorious for carrying divergent copies of 16S rRNA gene across distinct operons within a genome (Šorfová *et al*., 2008; Nováková *et al*., 2009, 2016), which seems to be the case also here. In *P. spumarius*, the presence of distinct sets of zOTUs of its gammaproteobacterial symbiont *Sodalis* in different individuals suggested different or divergent strains across populations(Kolasa *et al*., 2023); here most individuals of *M. laevis* had the same base set of zOTUs with some scattered genotypes for various populations. Unfortunately, so far, we lack information on the phylogenetic relationships among strains infecting different *Macrosteles* species: they might represent an ancient symbiont co-diverging with hosts, facultative endosymbiont capable of occasional horizontal transmission, result from independent acquisitions, or perhaps a combination of these possibilities. Likewise, so far, we lack genomic data on the nature of this unusually localised microbe. In other species, *Arsenophonus* has been established both as an obligate long-term endosymbiont or as a more recently acquired facultative, asin louse flies (Říhová *et al*., 2023). In the future, genomics and experiments should clarify both the relationships and roles of these common *Macrosteles* associates.

Other putative facultative endosymbionts - *Wolbachia*, *Rickettsia, Cardinium, and Lariskella* - were relatively rare in our collection, and patchily distributed across species and populations. *Wolbachia*, known for the breadth of its distribution across insect diversity, ability to manipulate host reproduction, and antiviral and other benefits it can confer (Weinert *et al*., 2015; Kaur *et al*., 2021) was found in only a few individuals. *Rickettsia* and *Cardinium* are also among the most widespread insect facultative endosymbionts, and have a range of effects comparable to *Wolbachia* (Weinert *et al*., 2015). We only found *Rickettsia* in *M. laevis* and *M. viridigreseus* but it has been reported previously in Japanese specimens of *M. sexnotatus* and *M. striifrons* (Ishii *et al*., 2013). *Cardinium* infected three *Macrosteles* species; while not reported from this genus before, it had been found in several species of planthoppers (Nakamura *et al*., 2009) and the leafhoppers *Scaphoideus titanus* (Marzorati *et al*., 2006) and *Euscelidius variegatus* (Chrostek *et al*., 2017). High infection prevalence in *M. cristatus*, *M. quadrilineatus* and *M. viridigreseus* suggests that at least in these species, it plays significant roles. A particularly interesting case is *Lariskella* in *M. maculosus*, to our knowledge the first record of this bacterium in Auchenorrhyncha. This Alphaproteobacterium has been reported from stink bugs, but also other arthropods such as fleas and ticks (Matsuura *et al*., 2012). Its fitness roles are unknown but in stinkbugs infected with the *Lariskella* no skewed sex ratio was observed, suggestive of their other functional roles, including the potential provision of nutrients (Matsuura *et al*., 2012).

Microbes impact insects in many ways, altering each other’s effects on host life history traits and biology (Sudakaran *et al*., 2017; Cornwallis *et al*., 2023). This includes reducing the transmissibility of vectored pathogens by other host-associated microbes. For example, in *Aedes aegypti* mosquitoes, artificial infection with *Wolbachia* largely blocks the transmission of dengue virus to humans (Edenborough *et al*., 2021). Due to the economic importance of *M. laevis* as a vector of plant-pathogenic phytoplasmas, assessing the relationship between the symbiont complement and the presence/abundance of *Phytoplasma* was among the primary original goals of this study. However, for *Arsenophonus*, the only microbe sufficiently prevalent to enable testing for *Phytoplasma*-protective effects, no significant association was found. It would be useful to systematically search for any such protective effects of other symbionts of diverse Auchenorrhyncha. However, our study has also revealed a limitation of the amplicon-sequencing search for infection: the limited detection sensitivity relative to the routine testing technique, nested PCR (Hafez *et al*., 2005). *Phytoplasma* abundance within host tissues varies among different stages of infection (Frost *et al*., 2011), and it can be assumed that at the time it is low, it may contribute only a small share of all 16S rRNA copies in a specimen. Depending on the amplicon sequencing depth, it may well avoid detection. Such screens could also be complicated by cross-contamination among samples that can happen on Illumina platforms during sequencing (Kircher *et al*., 2012). Likely, changes in the library preparation methods, including the optimization of primers or perhaps combining universal and specific primers, and the adjustment of sequencing depth, will make this approach much more reliable.

## Conclusion

We have shown that the simultaneous amplification and sequencing of host and microbial marker genes is a powerful technique for a broad and comprehensive survey of host-associated microbiota. Through sampling across species, as well as across populations and separate dates for one of the species, we aimed to simultaneously and broadly reconstruct spatio-temporal patterns of host mitochondrial diversity and microbial community composition, and use the former data to explain the latter. Such approaches, used repeatedly to characterise patterns in human microbiota (Ma *et al*., 2014; Gupta *et al*., 2017), have rarely been used for insects. We know of just two such approaches, one focused on aphid-parasitoid-symbiont networks (Ye *et al*., 2017) and the other, on the spittlebug *Philaenus spumarius* (Kolasa *et al*., 2023). But unlike the latter study, demonstrating a clear genetic structure among sampled insects and significant differences in microbiota composition among populations, we found little diversity and few clear patterns in *M. laevis*. There were no consistent differences among geographically distant populations or among sampling dates. In comparison to *P. spumarius*, or pea aphids that have also been systematically surveyed for multiple bacterial associates (Pilgrim *et al*., 2021), *M. laevis* microbiota are surprisingly stable. Future work will show whether stability is an unusual feature of this one species. On the other hand, the comparison of *M. laevis* with other *Macrosteles* species revealed significant differences and a much greater diversity of microbial associates, expanding our understanding of the microbiota of this widely distributed and ecologically significant genus. With potential methodological modifications that could enhance the detection success of pathogens such as *Phytoplasma*, high-throughput multi-target amplicon sequencing has the potential to greatly enhance our understanding of microbiome-related processes and patterns in natural populations and communities.

## Acknowledgments

We would like to thank Mike Wilson, Sam Mugford, and Adam Stroinski for contributing by providing us with important specimens. We are also greatly indebted to Mateusz Buczek and Marzena Marszałek for their technical skill and laboratory assistance.

## Data availability

The raw sequence has been deposited in the Sequence Read Archive of the National Centre for Biotechnology Information with the accession number PRJNA1044568

## References

Abram, P.K., Brodeur, J., Urbaneja, A., and Tena, A. (2019) Nonreproductive Effects of Insect Parasitoids on Their Hosts. Annu Rev Entomol 64: 259–276.

Apprill, A., McNally, S., Parsons, R., and Weber, L. (2015) Minor revision to V4 region SSU rRNA 806R gene primer greatly increases detection of SAR11 bacterioplankton. Aquat Microb Ecol 75: 129–137.

Avise, J.C. (2000) Phylogeography: The History and Formation of Species, Harvard University Press.

Bańkowska, R. (1989) Pipunculidae (Diptera) of moist meadows on the Mazovian lowland. memorabilia zool 43: 349–351.

Baumann, P. (2005) Biology of bacteriocyte-associated endosymbionts of plant sap-sucking insects. Annu Rev Microbiol 59: 155–189.

Bennett, G.M., Abbà, S., Kube, M., and Marzachì, C. (2016) Complete Genome Sequences of the Obligate Symbionts “ *Candidatus* Sulcia muelleri” and “ *Ca.* Nasuia deltocephalinicola” from the Pestiferous Leafhopper *Macrosteles quadripunctulatus* (Hemiptera: Cicadellidae). Genome Announc 4: e01604–15.

Bennett, G.M. and Moran, N.A. (2015) Heritable symbiosis: The advantages and perils of an evolutionary rabbit hole. Proc Natl Acad Sci USA 112: 10169–10176.

Bennett, G.M. and Moran, N.A. (2013) Small, Smaller, Smallest: The Origins and Evolution of Ancient Dual Symbioses in a Phloem-Feeding Insect. Genome Biology and Evolution 5: 1675–1688.

Campbell, M.A., Van Leuven, J.T., Meister, R.C., Carey, K.M., Simon, C., and McCutcheon, J.P. (2015) Genome expansion via lineage splitting and genome reduction in the cicada endosymbiont *Hodgkinia*. Proc Natl Acad Sci USA 112: 10192–10199.

Chac, L.D. and Thinh, B.B. (2023) Species Identification through DNA Barcoding and Its Applications: A Review. Biol Bull Russ Acad Sci 50: 1143–1156.

Chrostek, E., Pelz-Stelinski, K., Hurst, G.D.D., and Hughes, G.L. (2017) Horizontal Transmission of Intracellular Insect Symbionts via Plants. Front Microbiol 8: 2237.

Clements, J., Bradford, B.Z., Garcia, M., Piper, S., Huang, W., Zwolinska, A., et al. (2021) ‘Candidatus Phytoplasma asteris’ subgroups display distinct disease progression dynamics during the carrot growing season. PLoS ONE 16: e0239956.

Cornwallis, C.K., Van ’T Padje, A., Ellers, J., Klein, M., Jackson, R., Kiers, E.T., et al. (2023) Symbioses shape feeding niches and diversification across insects. Nat Ecol Evol 7: 1022–1044.

Deng, J., Bennett, G.M., Franco, D.C., Prus-Frankowska, M., Stroiński, A., Michalik, A., and Łukasik, P. (2023) Genome Comparison Reveals Inversions and Alternative Evolutionary History of Nutritional Endosymbionts in Planthoppers (Hemiptera: Fulgoromorpha). Genome Biology and Evolution 15: evad120.

Dicke, M., Cusumano, A., and Poelman, E.H. (2020) Microbial Symbionts of Parasitoids. Annu Rev Entomol 65: 171–190.

Douglas, A.E. (2015) Multiorganismal Insects: Diversity and Function of Resident Microorganisms. Annu Rev Entomol 60: 17–34.

Edenborough, K.M., Flores, H.A., Simmons, C.P., and Fraser, J.E. (2021) Using *Wolbachia* to Eliminate Dengue: Will the Virus Fight Back? J Virol 95: e02203–20.

Edgar, R.C., Haas, B.J., Clemente, J.C., Quince, C., and Knight, R. (2011) UCHIME improves sensitivity and speed of chimera detection. Bioinformatics 27: 2194–2200.

Elbrecht, V. and Leese, F. (2017) Validation and Development of COI Metabarcoding Primers for Freshwater Macroinvertebrate Bioassessment. Front Environ Sci 5:.

Engel, P. and Moran, N.A. (2013) The gut microbiota of insects – diversity in structure and function. FEMS Microbiol Rev 37: 699–735.

Fontaine, K.M., Cooley, J.R., and Simon, C. (2007) Evidence for Paternal Leakage in Hybrid Periodical Cicadas (Hemiptera: Magicicada spp.). PLoS ONE 2: e892.

Frey, J.E. and Frey, B. (2004) Origin of intra-individual variation in PCR-amplified mitochondrial cytochrome oxidase I of Thrips tabaci (Thysanoptera: Thripidae): mitochondrial heteroplasmy or nuclear integration?: Variation in PCR-amplified thrips COI sequences. Hereditas 140: 92–98.

Frost, K.E., Willis, D.K., and Groves, R.L. (2011) Detection and Variability of Aster Yellows Phytoplasma Titer in Its Insect Vector, Macrosteles quadrilineatus (Hemiptera: Cicadellidae). jnl econ entom 104: 1800–1815.

Gupta, V.K., Paul, S., and Dutta, C. (2017) Geography, Ethnicity or Subsistence-Specific Variations in Human Microbiome Composition and Diversity. Front Microbiol 8: 1162.

Hafez, H.M., Hauck, R., Lüschow, D., and McDougald, L. (2005) Comparison of the Specificity and Sensitivity of PCR, Nested PCR, and Real-Time PCR for the Diagnosis of Histomoniasis. Avian Diseases 49: 366–370.

Holzinger, W.E., Kammerlander, I., and Nickel, H. (2003) The Auchenorrhyncha of Central Europe. Die Zikaden Mitteleuropas, Volume 1: Fulgoromorpha, Cicadomorpha excl. Cicadellidae, BRILL.

Ishii, Y., Matsuura, Y., Kakizawa, S., Nikoh, N., and Fukatsu, T. (2013) Diversity of Bacterial Endosymbionts Associated with Macrosteles Leafhoppers Vectoring Phytopathogenic Phytoplasmas. Appl Environ Microbiol 79: 5013–5022.

Johnson, K.P., Dietrich, C.H., Friedrich, F., Beutel, R.G., Wipfler, B., Peters, R.S., et al. (2018) Phylogenomics and the evolution of hemipteroid insects. Proc Natl Acad Sci USA 115: 12775–12780.

Kaur, R., Shropshire, J.D., Cross, K.L., Leigh, B., Mansueto, A.J., Stewart, V., et al. (2021) Living in the endosymbiotic world of Wolbachia: A centennial review. Cell Host & Microbe 29: 879–893.

Kircher, M., Sawyer, S., and Meyer, M. (2012) Double indexing overcomes inaccuracies in multiplex sequencing on the Illumina platform. Nucleic Acids Research 40: e3–e3.

Kobiałka, M., Michalik, A., Walczak, M., Junkiert, Ł., and Szklarzewicz, T. (2016) Sulcia symbiont of the leafhopper Macrosteles laevis (Ribaut, 1927) (Insecta, Hemiptera, Cicadellidae: Deltocephalinae) harbors Arsenophonus bacteria. Protoplasma 253: 903–912.

Kolasa, M., Kajtoch, Ł., Michalik, A., Maryańska□Nadachowska, A., and Łukasik, P. (2023) Till evolution do us part: The diversity of symbiotic associations across populations of *Philaenus* spittlebugs. Environmental Microbiology 25: 2431–2446.

Kolde, R. (2019) pheatmap: Pretty Heatmaps.

Kumar, S., Stecher, G., Li, M., Knyaz, C., and Tamura, K. (2018) MEGA X: Molecular Evolutionary Genetics Analysis across Computing Platforms. Molecular Biology and Evolution 35: 1547–1549.

Leray, M., Ho, S.-L., Lin, I.-J., and Machida, R.J. (2018) MIDORI server: a webserver for taxonomic assignment of unknown metazoan mitochondrial-encoded sequences using a curated database. Bioinformatics 34: 3753–3754.

Ma, J., Coarfa, C., Qin, X., Bonnen, P.E., Milosavljevic, A., Versalovic, J., and Aagaard, K. (2014) mtDNA haplogroup and single nucleotide polymorphisms structure human microbiome communities. BMC Genomics 15: 257.

Mao, M. and Bennett, G.M. (2020) Symbiont replacements reset the co-evolutionary relationship between insects and their heritable bacteria. ISME J 14: 1384–1395.

Marzorati, M., Alma, A., Sacchi, L., Pajoro, M., Palermo, S., Brusetti, L., et al. (2006) A Novel *Bacteroidetes* Symbiont Is Localized in *Scaphoideus titanus*, the Insect Vector of Flavescence Dorée in *Vitis vinifera*. Appl Environ Microbiol 72: 1467–1475.

Matsuura, Y., Kikuchi, Y., Meng, X.Y., Koga, R., and Fukatsu, T. (2012) Novel Clade of Alphaproteobacterial Endosymbionts Associated with Stinkbugs and Other Arthropods. Appl Environ Microbiol 78: 4149–4156.

Matsuura, Y., Moriyama, M., Łukasik, P., Vanderpool, D., Tanahashi, M., Meng, X.-Y., et al. (2018) Recurrent symbiont recruitment from fungal parasites in cicadas. Proc Natl Acad Sci USA 115: E5970–E5979.

McCutcheon, J.P., Boyd, B.M., and Dale, C. (2019) The Life of an Insect Endosymbiont from the Cradle to the Grave. Current Biology 29: R485–R495.

McCutcheon, J.P., McDonald, B.R., and Moran, N.A. (2009) Convergent evolution of metabolic roles in bacterial co-symbionts of insects. Proc Natl Acad Sci USA 106: 15394–15399.

McFall-Ngai, M., Hadfield, M.G., Bosch, T.C.G., Carey, H.V., Domazet-Lošo, T., Douglas, A.E., et al. (2013) Animals in a bacterial world, a new imperative for the life sciences. Proc Natl Acad Sci USA 110: 3229–3236.

Meza-Lázaro, R.N., Poteaux, C., Bayona-Vásquez, N.J., Branstetter, M.G., and Zaldívar-Riverón, A. (2018) Extensive mitochondrial heteroplasmy in the neotropical ants of the *Ectatomma ruidum* complex (Formicidae: Ectatomminae). Mitochondrial DNA Part A 29: 1203–1214.

Michalik, A., Franco, D.C., Deng, J., Szklarzewicz, T., Stroiński, A., Kobiałka, M., and Łukasik, P. (2023) Variable organization of symbiont-containing tissue across planthoppers hosting different heritable endosymbionts. Front Physiol 14: 1135346.

Moran, N.A., McCutcheon, J.P., and Nakabachi, A. (2008) Genomics and Evolution of Heritable Bacterial Symbionts. Annu Rev Genet 42: 165–190.

Moran, N.A., Tran, P., and Gerardo, N.M. (2005) Symbiosis and Insect Diversification: an Ancient Symbiont of Sap-Feeding Insects from the Bacterial Phylum *Bacteroidetes*. Appl Environ Microbiol 71: 8802–8810.

Moussa, A., Guerrieri, E., Torcoli, S., Serina, F., Quaglino, F., and Mori, N. (2023) Identification of phytoplasmas associated with grapevine ‘bois noir’ and flavescence dorée in inter-row groundcover vegetation used for green manure in Franciacorta vineyards. J Plant Pathol 105: 1511–1519.

Murrell, P. (2014) gridBase: Integration of base and grid graphics.

Nakamura, Y., Kawai, S., Yukuhiro, F., Ito, S., Gotoh, T., Kisimoto, R., et al. (2009) Prevalence of *Cardinium* Bacteria in Planthoppers and Spider Mites and Taxonomic Revision of “ *Candidatus* Cardinium hertigii” Based on Detection of a New *Cardinium* Group from Biting Midges. Appl Environ Microbiol 75: 6757–6763.

Neuwirth, E. (2022) RColorBrewer: ColorBrewer Palettes.

Nováková, E., Hypša, V., and Moran, N.A. (2009) Arsenophonus, an emerging clade of intracellular symbionts with a broad host distribution. BMC Microbiol 9: 143.

Nováková, E., Hypša, V., Nguyen, P., Husník, F., and Darby, A.C. (2016) Genome sequence of Candidatus Arsenophonus lipopteni, the exclusive symbiont of a blood sucking fly Lipoptena cervi (Diptera: Hippoboscidae). Stand in Genomic Sci 11: 72.

Oliver, K.M., Degnan, P.H., Burke, G.R., and Moran, N.A. (2010) Facultative Symbionts in Aphids and the Horizontal Transfer of Ecologically Important Traits. Annu Rev Entomol 55: 247–266.

Parada, A.E., Needham, D.M., and Fuhrman, J.A. (2016) Every base matters: assessing small subunit rRNA primers for marine microbiomes with mock communities, time series and global field samples. Environmental Microbiology 18: 1403–1414.

Pilgrim, J., Thongprem, P., Davison, H.R., Siozios, S., Baylis, M., Zakharov, E.V., et al. (2021) Torix *Rickettsia* are widespread in arthropods and reflect a neglected symbiosis. GigaScience 10: giab021.

Qu, L.-Y., Lou, Y.-H., Fan, H.-W., Ye, Y.-X., Huang, H.-J., Hu, M.-Q., et al. (2013) Two endosymbiotic bacteria, Wolbachia and Arsenophonus, in the brown planthopper Nilaparvata lugens. Symbiosis 61: 47–53.

Quast, C., Pruesse, E., Yilmaz, P., Gerken, J., Schweer, T., Yarza, P., et al. (2012) The SILVA ribosomal RNA gene database project: improved data processing and web-based tools. Nucleic Acids Research 41: D590–D596.

Ramos, A., Esteves, M.B., Cortés, M.T.B., and Lopes, J.R.S. (2020) Maize Bushy Stunt Phytoplasma Favors Its Spread by Changing Host Preference of the Insect Vector. Insects 11: 600.

Říhová, J., Gupta, S., Darby, A.C., Nováková, E., and Hypša, V. (2023) Arsenophonus symbiosis with louse flies: multiple origins, coevolutionary dynamics, and metabolic significance.

Smith, A.H., Łukasik, P., O’Connor, M.P., Lee, A., Mayo, G., Drott, M.T., et al. (2015) Patterns, causes and consequences of defensive microbiome dynamics across multiple scales. Mol Ecol 24: 1135–1149.

Smith, A.H., O’Connor, M.P., Deal, B., Kotzer, C., Lee, A., Wagner, B., et al. (2021) Does getting defensive get you anywhere?—Seasonal balancing selection, temperature, and parasitoids shape real□world, protective endosymbiont dynamics in the pea aphid. Molecular Ecology 30: 2449–2472.

Songram, O., Sittipraneed, S., and Klinbunga, S. (2006) Mitochondrial DNA Diversity and Genetic Differentiation of the Honeybee (Apis cerana) in Thailand. Biochem Genet 44: 256–269.

Šorfová, P., Škeříková, A., and Hypša, V. (2008) An effect of 16S rRNA intercistronic variability on coevolutionary analysis in symbiotic bacteria: Molecular phylogeny of Arsenophonus triatominarum. Systematic and Applied Microbiology 31: 88–100.

Srivathsan, A., Ang, Y., Heraty, J.M., Hwang, W.S., Jusoh, W.F.A., Kutty, S.N., et al. (2023) Convergence of dominance and neglect in flying insect diversity. Nat Ecol Evol 7: 1012–1021.

Stecher, G., Tamura, K., and Kumar, S. (2020) Molecular Evolutionary Genetics Analysis (MEGA) for macOS. Molecular Biology and Evolution 37: 1237–1239.

Sudakaran, S., Kost, C., and Kaltenpoth, M. (2017) Symbiont Acquisition and Replacement as a Source of Ecological Innovation. Trends in Microbiology 25: 375–390.

Valdivia, C., Newton, J.A., Von Beeren, C., O’Donnell, S., Kronauer, D.J.C., Russell, J.A., and Łukasik, P. (2023) Microbial symbionts are shared between ants and their associated beetles. Environmental Microbiology 25: 3466–3483.

Walker, A.W., Martin, J.C., Scott, P., Parkhill, J., Flint, H.J., and Scott, K.P. (2015) 16S rRNA gene-based profiling of the human infant gut microbiota is strongly influenced by sample processing and PCR primer choice. Microbiome 3: 26.

Weinert, L.A., Araujo-Jnr, E.V., Ahmed, M.Z., and Welch, J.J. (2015) The incidence of bacterial endosymbionts in terrestrial arthropods. Proc R Soc B 282: 20150249.

Weintraub, P.G., Trivellone, V., and Krüger, K. (2019) The Biology and Ecology of Leafhopper Transmission of Phytoplasmas. In Phytoplasmas: Plant Pathogenic Bacteria - II. Bertaccini, A., Weintraub, P.G., Rao, G.P., and Mori, N. (eds). Singapore: Springer Singapore, pp. 27–51.

Wickham, H., Hester, J., Chang, W., and Bryan, J. (2022) devtools: Tools to Make Developing R Packages Easier.

Ye, Z., Vollhardt, I.M.G., Girtler, S., Wallinger, C., Tomanovic, Z., and Traugott, M. (2017) An effective molecular approach for assessing cereal aphid-parasitoid-endosymbiont networks. Sci Rep 7: 3138.

Zwolińska, A. and Borodynko□Filas, N. (2021) Intra and extragenomic variation between 16S rRNA genes found in 16SrI□B□ related phytopathogenic phytoplasma strains. Annals of Applied Biology 179: 368–381.

